# A novel model plant to study the light control of seed germination

**DOI:** 10.1101/470401

**Authors:** Zsuzsanna Mérai, Kai Graeber, Per Wilhelmsson, Kristian K. Ullrich, Waheed Arshad, Christopher Grosche, Danuše Tarkowská, Veronika Turečková, Miroslav Strnad, Stefan A. Rensing, Gerhard Leubner-Metzger, Ortrun Mittelsten Scheid

**Author notes:** To whom correspondence may be addressed., or, /9831. Current address: Max Planck Institute for Evolutionary Biology, Department of Evolutionary Genetics, 24306 Plön, Germany (Kristian K. Ullrich).

## Abstract

Timing of seed germination is crucial for seed plants and coordinated by internal and external cues, reflecting adaptations to different habitats. Physiological and molecular studies with lettuce and *Arabidopsis thaliana* have documented a strict requirement for light to initiate germination and identified many receptors, signalling cascades, and hormonal control elements. In contrast, seed germination of several other plants is inhibited by light, but the molecular basis of this converse response is unknown. We describe *Aethionema arabicum* (Brassicaceae) as a suitable model plant to investigate the mechanism of germination inhibition by light, as it comprises accessions with natural variation between light-sensitive and light-neutral responses. Inhibition is independent of light wavelength and increases with light intensity and duration. Gibberellins and abscisic acid are involved in the control of germination as in Arabidopsis, but transcriptome comparisons of light- and dark-exposed *Aethionema arabicum* seeds revealed that expression of genes for key regulators upon light exposure undergo converse changes, resulting in antipodal hormone regulation. This illustrates that similar modular components of a pathway in light-inhibited, light-neutral and light requiring germination among the Brassicaceae have been assembled by evolution to produce divergent pathways, likely as adaptive traits.

**Highlight:** In contrast to light requirement for Arabidopsis seed germination, germination of several *Aethionema arabicum* accessions is inhibited by light, due to antipodal transcriptional regulation of hormone balance.

## Introduction

Proper timing of germination is a critical step for the survival and propagation of seed plants. Light is a major environmental factor regulating seed germination, providing information about position in the soil, presence of competitors, day length and season. Plants living in various habitats have different optima for light conditions at the time of germination. Seeds can be categorized based on their response to white light during germination (Takaki, 2001). Light-requiring seeds (positive photoblastic) germinate only after a minimal exposure to light, while light-inhibited seeds germinate only in the dark. A third category of seeds is light-neutral, germinating both in light and darkness. The categories are not mutually exclusive: the germination of the ricegrass species *Oryzopsis miliacea* and the salt cress (*Thellungiella halophila*) is promoted by a short illumination but inhibited by continuous light (Koller and Negbi, 1959; Li *et al*., 2015; Negbi and Koller, 1964). The photoblastic classification considers only responses to the whole spectrum of white light, regardless of wavelength-specific effects. For example, germination of *Brachypodium* and other monocotyledonous species seeds is inhibited by blue light via cryptochrome receptors (Barrero *et al*., 2014), but induced by red light. In white light, the seeds germinate and therefore belong to the light-requiring seed category (Barrero *et al*., 2012). Seeds of many accessions from *Arabidopsis thaliana* and lettuce (*Lactuca sativa*), the model plants for research in this field, require a minimal light exposure for complete germination. Therefore, most insight into the role of light for seed germination originates from these light-requiring seed types (Casal and Sanchez, 1998; Shinomura *et al*., 1994; Shropshire *et al*., 1961). Only a limited number of plant species with light-inhibited seeds (negative photoblastic) have been described, for example *Phacelia tanacetifolia* (Chen, 1970; Chen, 1968) or *Citrullus lanatus*, for which seed germination is inhibited by the whole spectrum, including white, blue, red, and far-red light (Botha and Small, 1988; Thanos *et al*., 1991). The different photoblastic responses are likely an adaptive trait to harsh or quickly changing habitats: species with light-inhibited seeds often grow on sea coasts or in deserts where germination on the surface might be risky or deleterious. Light-inhibited germination might be advantageous to avoid direct sunlight, so that the germination occurs when the seeds are buried at various depths under shifting sand dunes (Koller, 1956; Lai *et al*., 2016; Thanos *et al*., 1991), although they are not strictly correlated with specific habitats (Vandelook *et al*., 2018).

In seeds of all photoblastic categories, light-regulated molecular changes during germination are associated with light perception through phytochromes regulating hormonal levels (Casal *et al*., 1998; Takaki, 2001). Gibberellic acid (GA) and abscisic acid (ABA) play a central role: GA induces germination and helps to break seed dormancy, while ABA is involved in dormancy establishment and maintenance. The balance of these two hormones determines seed fate (Finch-Savage and Leubner-Metzger, 2006). After light perception by PHYB, a cascade including several transcription factors and repressors leads to GA synthesis and ABA degradation in light-requiring seeds (Seo *et al*., 2009). A dual role of light has been shown in salt cress, where weak light promotes, but strong light inhibits GA accumulation (Li *et al*., 2015). In Arabidopsis, in red light, the expression of the GA biosynthesis genes *GA3 OXIDASE 1* (*AtGA3ox1*) and *GA3 OXIDASE 2* (*AtGA3ox2*) as well as the ABA-degrading *CYTOCHROME P450* gene family member *AtCYP707A2* are enhanced, whereas the ABA biosynthesis gene *NCED6* and the GA-degrading *GA2 OXIDASE 2* (*AtGA2ox2*) are repressed (reviewed in Seo *et al*., 2009; Shu *et al*., 2016). In contrast, knowledge about the molecular mechanism of light-inhibited and light-insensitive germination is missing, as no species with this seed trait has so far been established as a model for molecular and genetic approaches.

*Aethionema arabicum* (L.) Andrz. ex DC. (Brassicaceae) is an annual spring plant with a relatively small (203-240 Mbp), diploid genome that was recently sequenced (Franzke *et al*., 2011; Haudry *et al*., 2013). The Aethionemeae tribe, with approximately 57 species, is the earliest diverged tribe within the Brassicaceae and shares 70-80% genetic information with Arabidopsis. *Aethionema arabicum* (Aethionema in the following) is distributed in the Middle East and Eastern Mediterranean region. In this study, we show that the seeds of one accession from Turkey (TUR) germinate well in light, while the seeds of another accession from Cyprus (CYP) are strongly inhibited by the entire spectrum of visible light, in a quantitative manner. We characterize the physiological and molecular properties of seed germination in these two accessions and demonstrate that germination inhibition in light is associated with a decreased GA:ABA ratio, in contrast to the situation in light-requiring Arabidopsis. Transcriptome analysis revealed the involvement of similar regulatory components as in Arabidopsis but with opposite responses to light. In addition, we identified rich natural variation of the photoblastic phenotype within the Aethionemeae tribe. This makes Aethionema a very suitable model to investigate the variation in seed germination responses to light that exist in closely related species.

## Materials and Methods

### Plant material

Experiments were conducted with *Aethionema arabicum* (L.) Andrz. ex DC. accessions TUR ES1020 and CYP (obtained from Eric Schranz, Wageningen), Iran8456-1, Iran8456-2, and Iran8458 (obtained from Setareh Mohammadin, Wageningen), *Aethionema carneum* (Banks & Sol.) B.Fedtsch. accession KM2496, and *Aethionema heterocarpum* Trev. accessions KM2491 and KM2614 (obtained from Klaus Mummenhoff, Osnabrück). All seed material was produced by plants grown under 16 h light/19°C and 8 h dark/16°C diurnal cycles. After seed harvest, seed stocks were kept dry at 24°C for a minimum of 2 months.

### Germination test

All germination tests were conducted at the optimal temperature of 14°C in Petri dishes on 4-layer filter paper wetted with dH_2_O and supplemented with 0.1% plant preservative PPM (Plant Cell Technology). Germination assays were partly carried out with the addition of 10 µM GA_4+7_ (Duchefa), 10 µM fluridone, or 10 µM norflurazon (Sigma Aldrich and Duchefa) as indicated. All assays were done in three replicates with a minimum of 20 seeds each. Except for data in Fig. 1C, 1D and Fig. 2, white, red, and blue light exposure was uniformly set to 100 μmol m^-2^ s^-1^, light intensity and far-red exposure was set to 15 μmol m^-2^ s^-1^ for all experiments. For dark treatments, seeds were placed on wet filter paper under complete darkness. Diurnal and high light tests were carried out under an LED NS1 lamp with wide sun-like spectrum (Valoya). Light spectra and intensity were measured by LED Meter MK350S (UPRtek).

### Quantitative RT-PCR

Before imbibition, seeds were sterilized with chlorine gas for 10 minutes. After 23 hours incubation in dark or light, seeds with intact seed coats were collected for RNA extraction, three biological replicates for each sample. RNA was extracted as described (Oñate-Sánchez and Vicente-Carbajosa, 2008). Three µg total RNA was treated with DNase I (Thermo Scientific) and precipitated with 1/10 volume of 3 M sodium acetate (pH 5.2) and 2.5 volume of ethanol. One µg DNase I-treated RNA was used for cDNA synthesis with random hexamer primers using the RevertAid H Minus First Strand cDNA Synthesis kit (Thermo Scientific). qRT-PCR reactions were performed in a Lightcycler^®^ 96 System (Roche) in duplicate, using FastStart Essential DNA Green Master mix (Roche) and primer pairs listed in Table S2, with the following parameters: 95°C for 10 min, 45 cycles with 95°C for 10 sec, and 60°C for 30 sec, and 1 cycle with 95°C for 10 sec, 60°C for 30 sec, and 97°C for 1 sec to obtain the melting curve for each reaction. Ct values were calculated using Lightcycler^®^ 96 Software (Roche). The geometric mean of Aethionema orthologues of ACTIN2 (*AearACT2*, AA26G00546), POLYUBIQUITIN10 (*AearUBQ10*, AA6G00219), and ANAPHASE-PROMOTING COMPLEX2 (*AearAPC2*, AA61G00327) was used for normalization. For each gene, the expression level under white light is presented as fold change relative to the level of the dark samples where the average expression was set to one.

### *Aethionema arabicum* genome and annotations

Genome scaffolds and accompanying gene feature format (GFF) file of *Aethionema arabicum* genome version 2.5 was obtained from (Haudry *et al*., 2013). The gene models were searched against the non-redundant database of NCBI (release 13-05-2015) using BLAST (Altschul *et al*., 1990). GO-terms were retrieved using BLAST2GO version 2.5 (Conesa *et al*., 2005) along with best hit and its description. The coding DNA sequence (CDS) of each gene model was translated into amino acid sequence using R package biostrings version 2.32.0 (https://www.bioconductor.org/packages/release/bioc/html/Biostrings.html) and then blasted against the uniprot database (release 2015_10) and The Arabidopsis Information Resource (TAIR, TAIR10_pep_20110103_representative_gene_model_updated) for extracting the best hit and its description. Results were filtered for having at least 80% query coverage, according to (Rost, 1999), to unambiguously detect homologous sequences. Sequence data from *Aethionema arabicum* can be found in CoGe database (https://genomevolution.org/coge/) under the following gid: v2.5, id33968. Accession numbers used in this study are listed on Supplementary Table S3.

Description of RNA-seq sample preparation, RNA-seq data handling, phylogenetic analysis and measurement of hormone levels can be found in Supplementary Methods S1.

## Results

### Light inhibits seed germination in *Aethionema arabicum*

The Aethionema accession TUR originates from Turkey, Konya (accession ES1020) and was used to generate the reference genome (Haudry *et al*., 2013) and to characterize its interesting seed and fruit dimorphism (Lenser *et al*., 2016). The Aethionema accession CYP comes from the Kato-Moni region in Cyprus (Mohammadin *et al*., 2017b). Both accessions were propagated for several generations under the same conditions in a growth chamber (Fig. 1A). Seeds of both accessions germinate optimally at 14°C, and all experiments were performed at that temperature. Testing the light dependence, we found that TUR seeds germinate under white light or in darkness. CYP seeds germinate well in darkness but are strongly inhibited under white light (Fig. **1**B). Species with light-requiring seeds need various periods of illumination for germination induction, ranging from seconds to days (Bewley *et al*., 2013). Reversely, light inhibition of germination can be exerted with a wide range of photon irradiance, from a relatively weak ~17 µmol m^-2^ s^-1^ light intensity in *Citrullus lanatus* (Botha and Small, 1988) to strong irradiance with 1000 µmol m^-2^ s^-1^ in some desert plants (Botha and Small, 1988; Lai *et al*., 2016). Therefore, we tested the germination of TUR and CYP seeds with different light intensities ranging from 0.1 to 350 µmol m^-2^ s^-1^ (Fig. **1**C). Increased light intensity gradually decreased the germination of CYP seeds: at 120 µmol m^-2^ s^-1^ only ~10% of the seeds germinated, and at 350 µmol m^-2^ s^-1^ the inhibition was complete. Germination of TUR seeds was light-neutral in this range, with only a slight reduction at 350 µmol m^-2^ s^-1^ (Fig. **1**C). As the irradiance can be much stronger at the geographic origin of the species, we further tested the germination of TUR seeds under 1600 µmol m^-2^ s^-1^ white light, with a wide sun-like spectrum. Although the germination of TUR seeds was slower, almost all seeds germinated under strong light, therefore we categorized TUR seeds as neutral to light (Fig. **1**D).

**Fig. 1.**
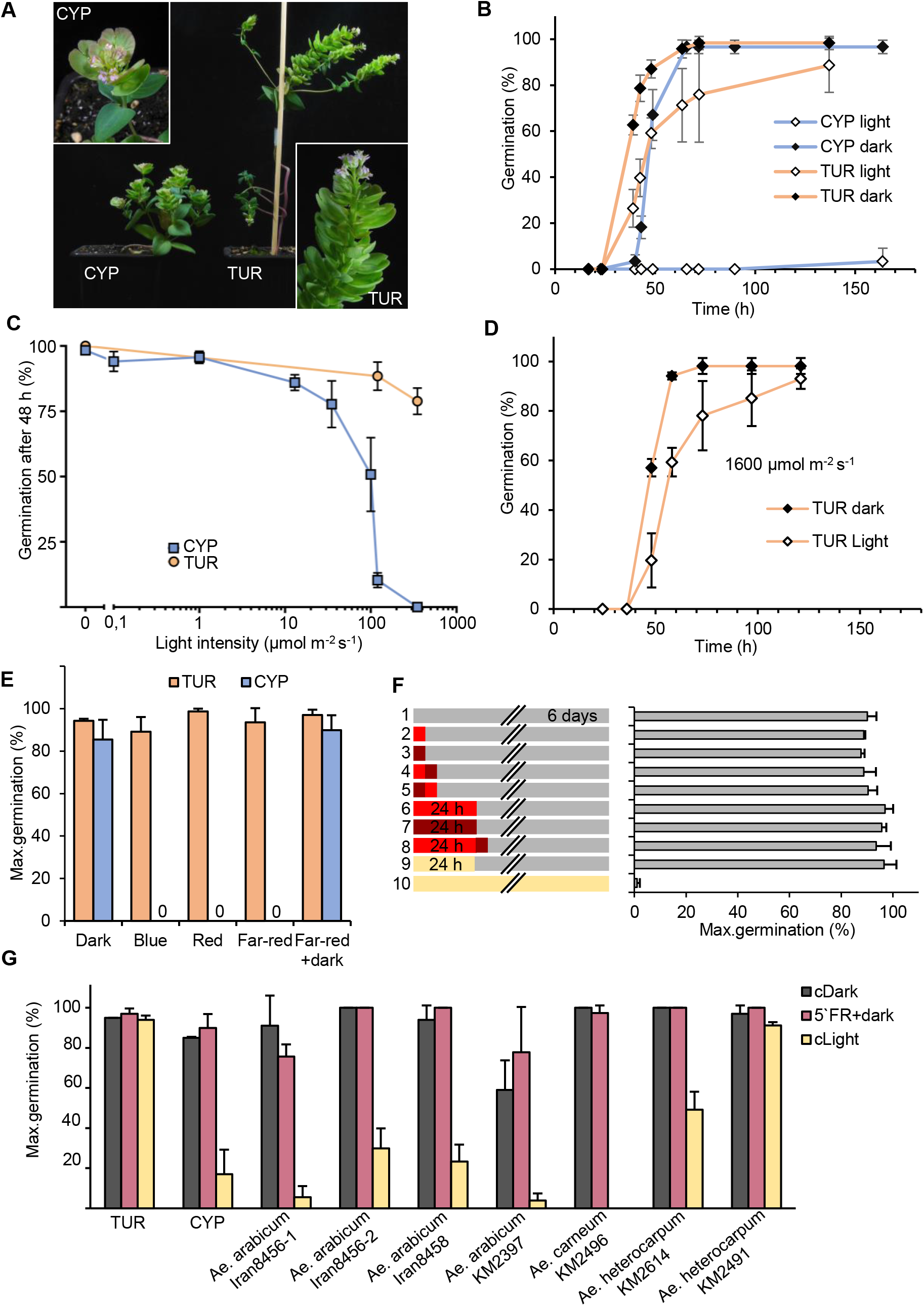
Light inhibition of seed germination in *Aethionema arabicum* accession CYP. **(A)** Eight-week-old *Aethionema arabicum* TUR (Turkey) and CYP (Cyprus) plants grown in a growth chamber. **(B)** Percentage of germinating seeds kept in darkness or under 100 μmol m-^2^ s^-1^ white light, scored over time. **(C)** Percentage of germinating seeds kept at various light intensities: 0.1, 1, 13, 35, 100, 120 and 350 μmol m^-2^ s^-1^ white light or in darkness (indicated as 0 μmol m^-2^ s^-1^), scored after 2 d. (**D**) Percentage of germinating TUR seeds kept in darkness or under 1600 μmol m^-2^ s^-1^ white light, scored over time. **(E)** Percentage of germinating seeds in continuous white, red, blue, or far-red light, dark, or 5 min far-red light followed by darkness, scored after 6 d. **(F)** Percentage of germinating CYP seeds after various light treatments. Left panel indicates the treatment, right panel indicates the percentage of germinating seeds, scored after 6 d. Red, dark red and cream rectangles indicate red, far-red or white light exposure, respectively. Grey bars indicate dark periods. Short and long red/farred rectangles indicate 5 min and 24 h exposure, respectively. **(G)** Percentage of germinating seeds from different *Ae. arabicum* accessions and closely related species, kept in dark (black columns), light (yellow columns), or in dark after a 5 min far-red pulse at imbibition (red columns), scored after 6 d. Error bars represent standard deviation from three independent replicates.

Light-dependent germination often requires exposure to a specific part of the light spectrum. Therefore, we tested if the inhibition of CYP seed germination depended on the wavelength. Continuous monochromatic blue, red, and far-red light were equally effective at inhibiting the germination of CYP seeds (Fig. **1**E), while TUR seeds could germinate under any light condition, including continuous far-red light that inhibits phytochrome-mediated light-induced germination in Arabidopsis and lettuce (Borthwick *et al*., 1952; Shropshire *et al*., 1961). Importantly, short (5 min) or longer (24 h) of either far-red or red illumination at the time of imbibition, followed by 6 days in darkness, allowed germination of CYP seeds equally well. This indicates that (i) the induction of germination is independent of the active form of phyB and ii) the light inhibition is not established in this time scale (Fig.**1E, F**).

TUR and CYP accessions cluster closely together in a network analysis of several *Aethionema arabicum* accessions (Mohammadin *et al*., 2017b). To determine if light inhibition of CYP seed germination is a unique trait in the genus Aethionema, we investigated the germination phenotype of other available accessions including the closest relatives, *Ae. heterocarpum* and *Ae. carneum* (Lenser *et al*., 2016, and Supplementary Table 1; Mohammadin *et al*., 2017b), after propagating the seeds under the same controlled conditions as for TUR and CYP. None of the tested accessions required light, as all of them germinated similarly well in constant darkness or after a five minutes far-red pulse followed by darkness (Fig. 1G). Under white light, we observed variations in the response: seed germination of *Ae. carneum* and two *Ae arabicum* (Iran 8456-1 and KM2397) accessions was clearly light-inhibited, one *Ae. heterocarpum* accession (KM2491) had light-neutral seeds, while the germination of 3 accessions (*Ae. arabicum* Iran 8458, Iran 8456-2 and *Ae. heterocarpum* KM2614) was partially inhibited by light (Fig. 1G). This reveals natural variation for the negative or neutral photoblastic phenotype, suggesting that light-inhibited or light-neutral seed germination may be an adaptive trait in the Aethionemeae tribe, and provides interesting material for genetic analysis of the phenomenon.

### Diurnal regulation of CYP seed germination

To better understand the ecological relevance of light-inhibited germination of CYP seeds, we also tested germination under different diurnal conditions. Again, TUR seeds were unaffected and germinated well under any daylength. Interestingly, at a lower range of light intensity, the CYP seeds germinated well under short day conditions (cycles of 8 h light/16 h darkness, LD8/16), compared to LD12/12 with partial inhibition or LD16/8 cycles with complete inhibition (Fig. 2A). Remarkably, at higher light intensity similar to conditions in the natural habitat (1600 µmol m^-2^ s^-1^), still ~40% of the CYP seeds germinated under short day (Fig. 2B). Gradual inhibition was also observed with hourly alternating light exposure (Fig. 2C). However, longer uninterrupted light periods resulted in stronger inhibition than more frequent alterations, despite equal daily fluence in LD12h/12h compared to LD30min/30min (Fig. 2A and Fig. 2C). These data indicate that CYP seeds integrate regime, duration, and intensity of illumination into their germination regulation.

**Fig. 2.**
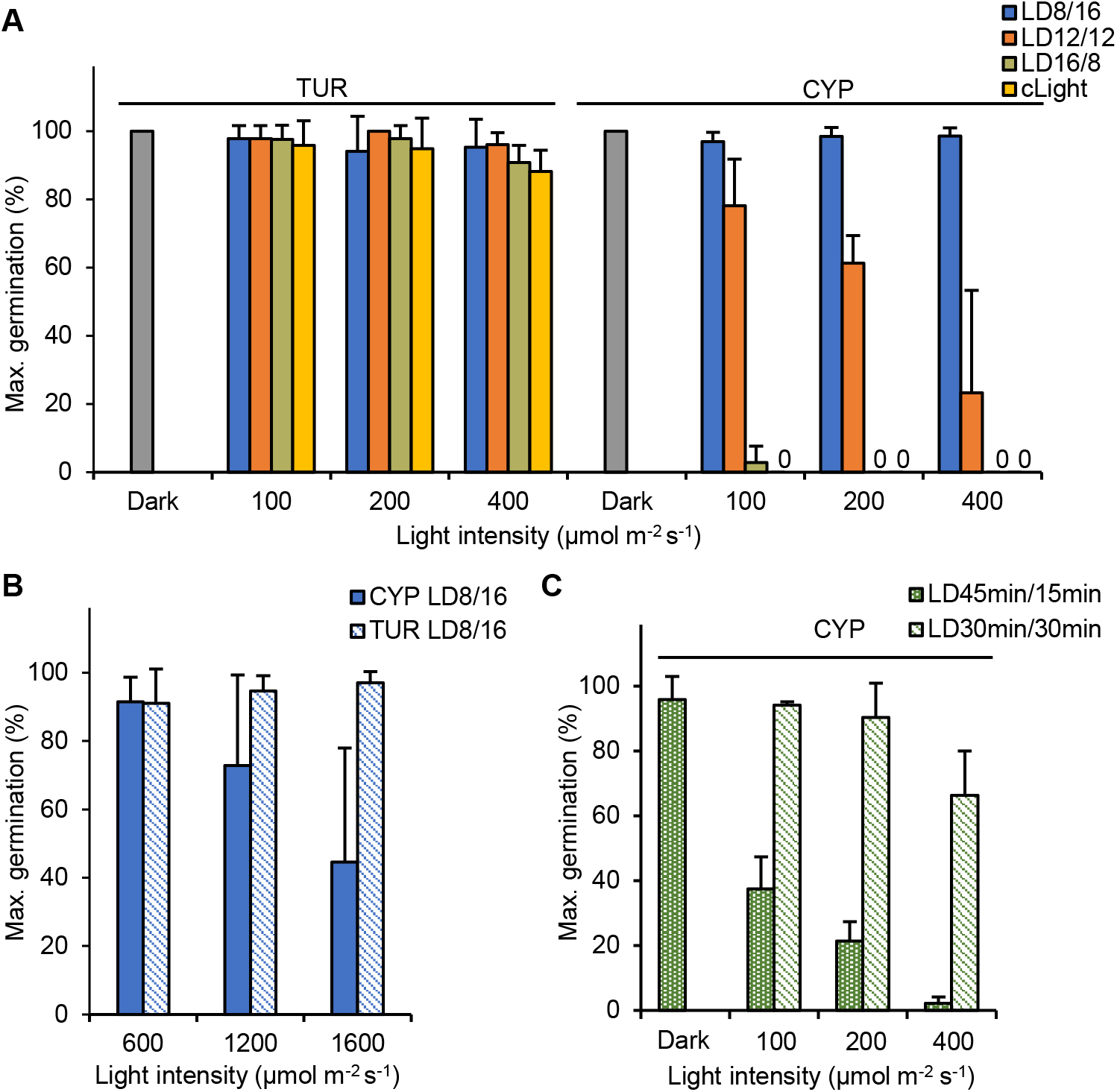
Diurnal regulation of CYP seed germination. **(A-B)** Percentage of germinating seeds kept under different diurnal regimes, scored after 6 d. LD8/16: 8 h light and 16 h dark, LD12/12: 12 h light and 12 h dark, LD16/8: 16 h light and 8 h dark. In addition, seeds were kept under continuous dark (Dark) or light (cLight). 0 indicates zero germination. **(C)** Percentage of germinating CYP seeds under continuously repeated 30 min light/30 min dark cycles (LD30min/30min) or 45 min light and 15 min dark cycles (LD45min/45min), scored after 6 d. Error bars represent standard deviation from three independent replicates.

### Light-neutral and light-inhibited seeds differ in their transcriptomes

To better understand the light inhibited seed germination phenotype in Aethionema, we performed transcriptome analysis of the TUR and CYP accessions. Seeds were imbibed and kept in darkness (D) or under 100 µmol m^-2^ s^-1^ white light (WL) for 23 hours, which was determined as the start point for the completion of germination (~1% seeds in the responsive populations with emerged radicles). It is important to note that CYP seeds are fully able to germinate if transferred to darkness after 23 h of light exposure, therefore not dormant (Fig. 1F). Only seeds with intact seed coats were sampled and used for the preparation of RNA libraries. Comparison of dark- and light-exposed samples revealed 168 differentially expressed genes in the TUR accession: 51 genes upregulated and 117 genes downregulated in seeds kept in darkness compared to those in light (Fig. 3A, Dataset **S1**). In the CYP accession, we found 214 differentially regulated genes, 105 up- and 109 down-regulated, in seeds in darkness (Fig. 3A, Dataset **S2**). Considering the close relation between the accessions, the overlap of 93 genes commonly differentially regulated in TUR and CYP was surprisingly small, while 75 and 121 genes were light-dependent only in TUR or CYP, respectively (Fig. 3A). This comparison, however, did not consider the genetic differences between the two accessions. We therefore compared the transcriptome of each condition between the TUR and CYP samples. In seeds kept in darkness, 564 genes were differentially expressed between the two accessions (Fig.**3B**, Dataset **S3**). The number of differentially expressed genes between TUR and CYP light-exposed samples was much higher (969), matching the expectation for a larger difference upon light exposure and a different physiological state regarding the capability for germination. Among the 969 genes, 613 were expressed at higher levels in TUR while 356 gene were more expressed in CYP (Fig. 3B, Dataset **S4**). Nearly half of the genes (469) were found in the overlap, indicating transcriptional differences between the two genotypes independently from the light conditions. However, these genes might undergo further light regulation that could contribute to the phenotypes. To distinguish these possibilities, we hypothesized that the light-inhibited germination in CYP should be associated with genes that are (i) light-regulated in CYP seeds and (ii) differentially expressed in light-exposed TUR and CYP seeds. This selection would include genes that are possibly also light-regulated in TUR seeds to a lesser extent than in CYP seeds, or their induction/repression would be similar in both accessions but their absolute level results in different expression upon light induction. Both criteria were fulfilled for 87 genes (Fig. **3**C, D, Dataset **S5**, Fig. **S1**). Out of the 87 genes, 16 were upregulated and 66 were downregulated in the light-exposed CYP seeds (Fig. 3D). Surprisingly, with only 3 exceptions, the direction of light-regulated transcript level changes was the same in both accessions, although the response was much more pronounced in CYP compared to TUR (Fig. 3D, **S1**). For 85 out of 87 genes we could identify the Arabidopsis orthologues (Dataset **S5**). Based on the TAIR10 database description, 10 of these genes are linked to light stimuli and 11 to cell wall organization or biogenesis (Fig. 3E). Additionally, 10 genes are involved in hormonal signaling or response (Fig. 3E). Genes associated with ABA biosynthesis or degradation were not present among these genes (Fig. 3E, Dataset **S5**). However, the most strongly upregulated transcript (>50-fold induction in CYP WL seeds) was *AA18G00108*, encoding a gibberellin 2-oxidase. Phylogenetic analysis and synteny (Fig. **S2**) of the genomic position confirmed that Aethionema *AA18G00108* is the orthologue of *AtGA2ox3* (*AT2G34555*), encoding a protein with C-19 gibberellin 2-beta-dioxygenase activity and involved in the degradation of GA, therefore expected to negatively influence germination.

**Fig. 3.**
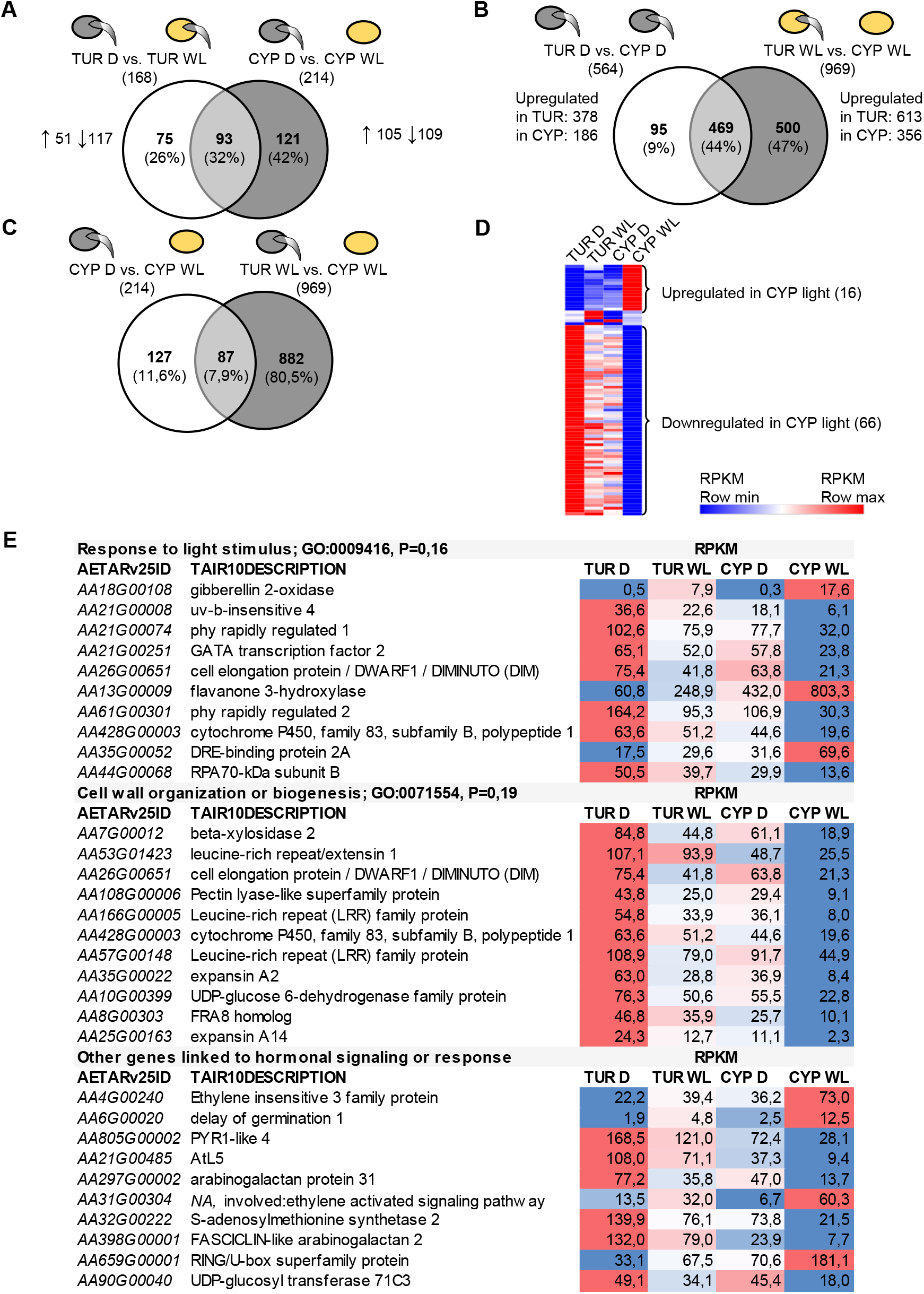
Transcriptome analysis of Aethionema TUR and CYP seeds by RNA-seq. **(A-C)** Number of differentially expressed genes and Venn diagrams showing the proportion of overlapping genes in pairwise comparisons between TUR (Turkey accession) and CYP (Cyprus accession), D (dark, grey seed icon), or WL (white light-exposed seeds, yellow icon). Icons with germinating seeds indicate the capability of samples for germination, noting that the seeds did not yet germinate at the timepoint of sampling. **(D)** Heatmap indicating the relative expression of the overlapping 87 genes in the four treatments. Coloring is based on the RPKM values. A detailed list is in Supplemental Fig. 1. **(E)** Gene list of three selected GO terms with average RPKM values and the relative expression differences indicated by shade of blue (lowest RPKM value) or red (highest RPKM value).

### Transcriptome analysis reveals differences in light-mediated hormonal responses

In most species, the balance of GA and ABA hormones is an important component of the light-regulated germination. We therefore tested individual transcript level changes of genes involved in GA and ABA biosynthesis in D- and WL-germinated TUR and CYP seeds. Overall, upon light exposure in Aethionema seeds, we found slightly increased transcript levels of the genes encoding ABA biosynthetic enzymes and a decrease of the main GA biosynthetic enzymes (Fig. **4**A, B). Expression of the Aethionema orthologues of *ABA1*, *ABA2*, and *ABA3* was slightly but significantly upregulated upon light exposure, both in the light-neutral TUR and the light-inhibited CYP seeds (Fig. 4A). *NCED6* and *NCED9*, encoding the 9-cis epoxycarotenoid dioxygenase gating the catalysis of 9`-cis neoxanthin, the rate-limiting step of ABA synthesis, are known to be transcriptionally downregulated in Arabidopsis upon red light induction (Oh *et al*., 2007; Seo *et al*., 2006; Seo *et al*., 2009). In both Aethionema accessions, *AearNCED5* was significantly upregulated in light, while the level of *AearNCED9* was similar in dark and light (Fig. 4A). In contrast, the expression of the gene for the ABA-deactivating enzyme *AearCYP7072A* was also elevated, in this case matching the observations in Arabidopsis seeds (Fig. 4A) (Oh *et al*., 2007; Seo *et al*., 2006). However, the light-induced transcript level changes of ABA-related genes did not explain the difference between the TUR and CYP seeds, as the direction and the intensity of changes were similar in the two accessions.

**Fig. 4.**
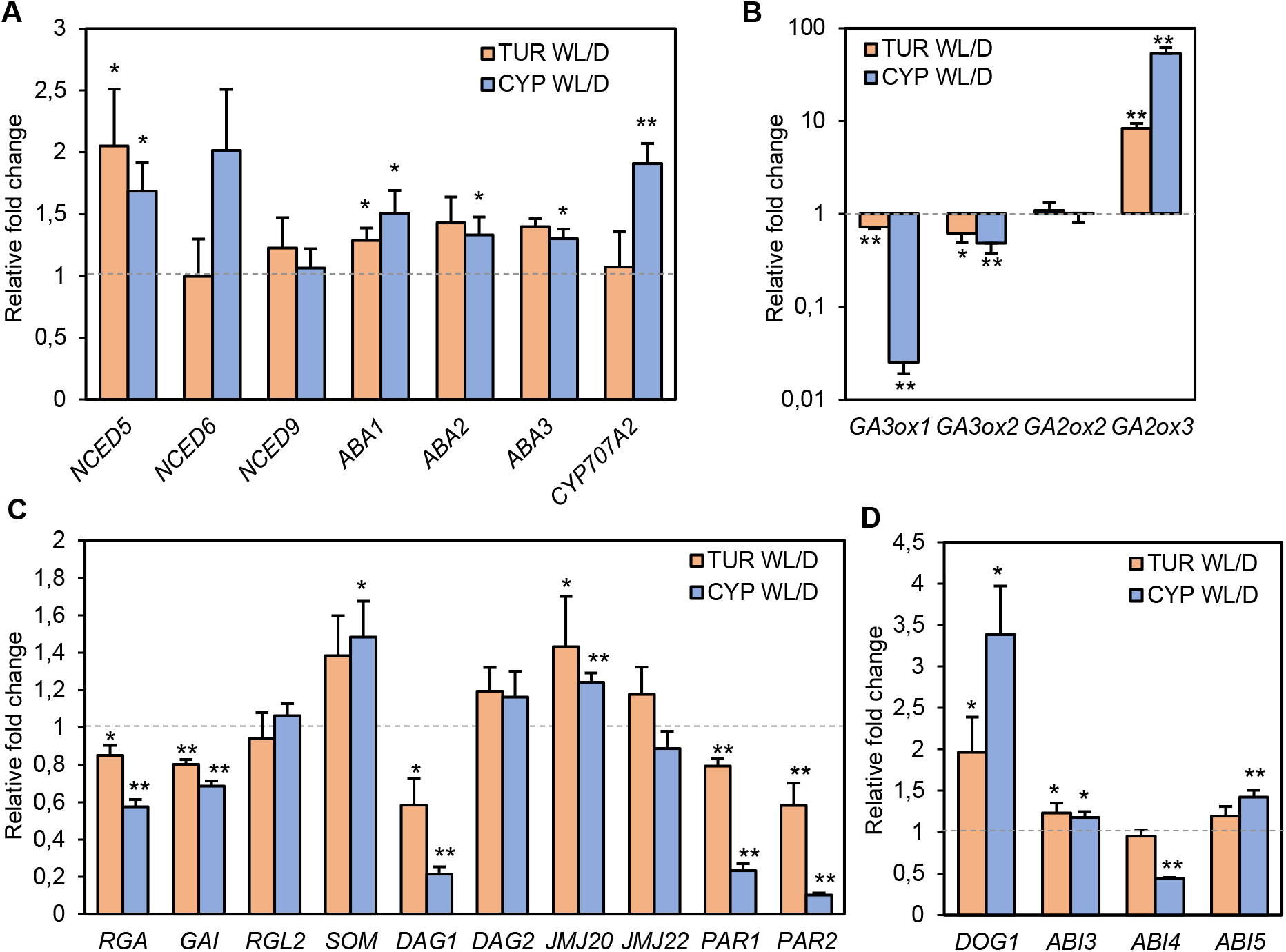
Analysis of the Aethionema light-regulated transcriptome network. **(A-D**) Quantitative RT-PCR for selected genes. The expression level under white light (WL) is presented as fold change relative to the average expression level in darkness (D) set to 1 (indicated with grey dashed line). Significance was tested using the Student T-test. * p< 0.1 or ** p<0.05 indicate if the white-light and dark expression level of a gene is significantly different. Error bars represent standard deviation from three independent biological replicates. TUR (Turkey accession), CYP (Cyprus accession), see text for gene acronyms.

Light-induced GA accumulation in Arabidopsis seeds is mediated by the enhanced expression of GA biosynthetic enzymes, encoded by *AtGA3ox1* and *AtGA3ox2*, and the decrease of the GA-deactivating GA2-oxidase encoded by *AtGA2ox2* (Seo *et al*., 2006; Yamaguchi *et al*., 1998). We found a reciprocal situation in Aethionema seeds: the expression of both *AearGA3ox1* and *AearGA3ox2* was decreased (Fig. 4B). In good association with the RNA-seq results, the expression of *AearGA2ox3* upon WL illumination was increased in both accessions (Fig. 4B). Remarkably, this activation of *AearGA2ox3*, as well as the repression of *AearGA3ox1* were considerably more pronounced in the light-inhibited CYP seeds compared to TUR seeds (Fig. 4B). This validates the previous observation that the direction of light-regulated transcript level changes concurs in both accessions but is much more pronounced in CYP seeds (Fig. 3D, 4B). It also suggested that the TUR and CYP accessions might differ in the regulatory network upstream of GA genes. Therefore, we further investigated the transcriptome for regulation by the known factors involved in light-mediated hormonal responses in seeds.

Upon light reception in Arabidopsis, the active P_fr_ form of phytochrome B interacts with PIL5, a basic helix-loop-helix protein, facilitating its degradation by the 26S proteasome (Oh *et al*., 2004; Oh *et al*., 2007; Shen *et al*., 2008). Under dark or FR conditions, PIL5 mediates the stable expression of *GAI* and *RGA*, two genes encoding DELLA proteins which are negative components of GA signaling (Oh *et al*., 2007; Piskurewicz *et al*., 2009). In parallel, PIL5 directly stimulates the expression of *SOM* (*SOMNUS*), a negative regulator of light-dependent germination that controls the expression of ABA and GA metabolic genes (Kim *et al*., 2008). Both *pil5* and *som* mutants of Arabidopsis germinate in a light-insensitive manner (Kim *et al*., 2008; Oh *et al*., 2004). The link between SOM and the *GA3ox1*/2 genes is formed by two jumonji-domain proteins, JMJ20 and JMJ22, that are directly repressed by SOM and support germination by removing the repressing histone 4 arginine 3 methylation from *GA3ox1*/2, allowing their expression (Cho *et al*., 2012). Another negative regulator of seed germination is DAG1 (DOF AFFECTING GERMINATION1) that is under indirect positive control downstream of PIL5 and directly represses the transcription of *GA3ox1* and *DAG2*, which was recently identified as a positive regulator of light-induced germination (Boccaccini *et al*., 2014; Gabriele *et al*., 2010; Santopolo *et al*., 2015). However, we found only moderate transcript level changes (fold changes between 0.5-1.5*) of the Aethionema orthologues of *RGA*, *GAI*, *RGL2*, *SOM*, *DAG2*, *JMJ20*, and *JMJ22* in light-exposed seeds compared to those in dark (Fig. 4C). The expression of *DAG1* was decreased in both accessions in the light-exposed seeds, similar to its response in Arabidopsis (Fig. 4C). *PHYTOCHROME RAPIDLY REGULATED 1* and *2* (*PAR1* and *PAR2*) were both downregulated in WL seeds (Fig. 3E, Dataset **S5**). In Arabidopsis, *PAR1* and *2* are negative factors in shade avoidance (Roig-Villanova *et al*., 2007) and promote seedling de-etiolation under different light conditions, likely through interaction with PIF proteins (Zhou *et al*., 2014). Their role during seed germination has not been elucidated, although available transcriptome data suggests that *PAR2* is repressed by PIL5 and upregulated in red light in seeds (Shi *et al*., 2013). Importantly, we also confirmed the RNA-seq results that *AearPAR1* and *AearPAR2* are downregulated in WL seeds (Fig. 3E, 4C, *AA21G00074* and *AA61G00301*).

Germination under unfavorable conditions can be avoided by the establishment of seed dormancy, strongly correlated with the key dormancy protein DOG1 (DELAY OF GERMINATION 1) (Bentsink *et al*., 2010; Bentsink *et al*., 2006; Footitt *et al*., 2011; Graeber *et al*., 2014; Kerdaffrec *et al*., 2016). The expression of the *DOG1* gene is regulated by environmental signals and highly variable among Arabidopsis accessions (Chiang *et al*., 2011; Finch-Savage and Footitt, 2017; Kendall *et al*., 2011). Therefore, we tested if the light exposure of TUR or CYP seeds enhances *DOG1* expression. *AearDOG1* was indeed found among the light-responsive genes upregulated in CYP (Fig. 3E, AA6G00020). qRT-PCR data confirmed that *DOG1* expression was significantly enhanced in both accessions in light-exposed seeds, but the increase was more pronounced in the light-inhibited CYP seeds (Fig. 3D). These results indicate that light can indeed influence the dormancy level in responsive seeds. The expression of Aethionema orthologues of ABA-responsive transcription factors linked to dormancy, including *ABI3*, *ABI4*, and *ABI5* (*ABA INSENSITIVE 3*, *4*, *5*) (Dekkers *et al*., 2016; Shu *et al*., 2013) showed significant, but only moderate differences in light- and dark- exposed seeds (Fig. 3D). Taken together, (i) Aethionema uses similar key regulatory components to control germination as Arabidopsis, (ii) but the direction of several transcript level changes in the light-exposed seeds are opposite in seeds of both Aethionema accessions compared to Arabidopsis, and (iii) at least two genes related to GA biosynthesis or degradation (*GA3ox1* and *GA2ox3*) show a much more pronounced response in the light-inhibited CYP seeds compared to the light-neutral TUR seeds and might be responsible for the observed contrasting effect of light on germination.

### The GA:ABA ratio decreases during light inhibition of Aethionema seed germination

To test if the differential expression of the GA- or ABA-related genes would indeed affect the levels of the respective bioactive hormones, we compared GA and ABA levels in TUR and CYP seeds during the different germination schemes. For germination induction of Arabidopsis seeds, the most active form of GA is GA_4,_ among other GAs studied (Derkx *et al*., 1994). The absolute GA_4_ hormone level was similar in both Aethionema accessions in seeds kept in darkness and in light-exposed TUR (Fig. 5A). In contrast, and in good correlation with the light-induced repression of *GA3ox1* gene expression (Fig. 4B), we observed a significant decrease of the bioactive GA_4_ in CYP seeds kept under light for 23 hours (Fig. 5A, **S3**). Moreover, the level of GA_9_, the biosynthetic precursor of GA_4,_ was also significantly higher in light-exposed CYP seeds compared to TUR, indicating that the GA_9_ → GA_4_ conversion is less efficient in CYP seeds under light (Fig. 5A). According to the enhanced expression of the catabolic *GA2ox3* in CYP light-exposed seeds, we expected an increased level of the catabolic GA_34_ in CYP seeds. Although we could not observe this tendency during the 23-hour period, it might be due to the slower turnover of GA (Fig. 5A). Absolute ABA levels were higher in CYP seeds under both regimes, in agreement with the later onset of CYP germination in the dark compared to TUR, and with significantly higher levels in light-exposed CYP seeds compared to TUR (Fig. 5B). As seed germination is determined by the balance of the two antagonizing hormones, we calculated the molar ratio of GA and ABA. Interestingly, the GA:ABA ratio decreased in both accessions under light (Fig. 5C), but to a much larger extent in light-exposed CYP seeds, which had the lowest GA:ABA ratio of all. This indicates a threshold for the hormonal control below which germination of CYP seeds under continuous light exposure is not possible.

**Fig. 5.**
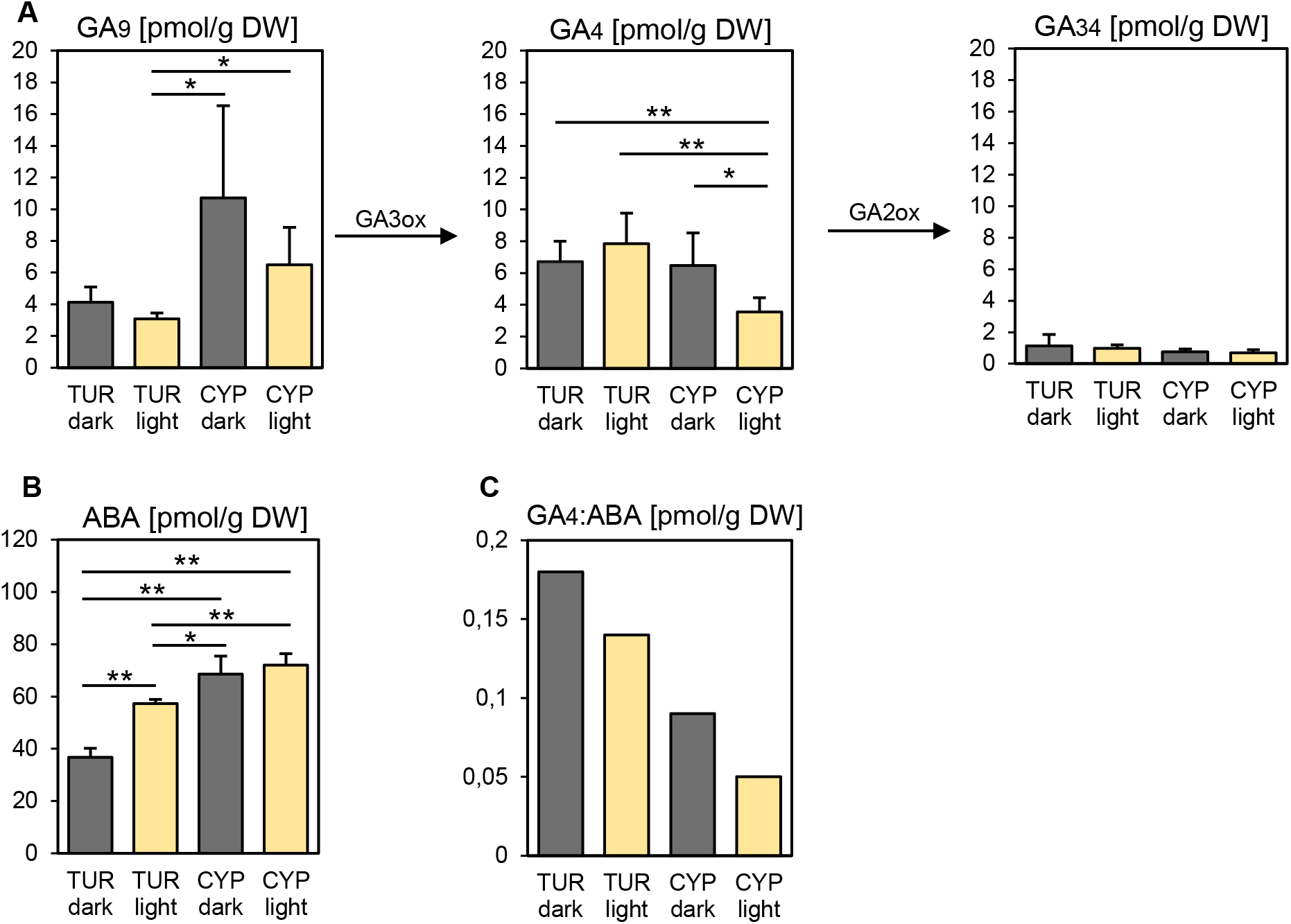
Decreased GA:ABA hormone ratio in light-exposed Aethionema CYP seeds. Hormone concentrations (in pmol per gram dry weight (DW)) of GA_9_ precursor, bioactive GA_4_ and GA_34_ catabolite **(A)** and ABA **(B)** are shown for dark- (grey columns) and light- (cream columns) exposed TUR and CYP seeds. **(C)** Molar ratio of GA_4_ and ABA. Significance was tested using the Student T-test, * p< 0.1, ** p<0.05 indicate significant differences. Error bars represent standard deviation between five independent replicates. Values for other GA metabolites are presented in Supplemental Fig. 3.

### GA and ABA are involved in light inhibition of CYP seed germination

In Arabidopsis, germination of far-red-exposed seeds can be rescued by removal of the seed coat and the endosperm layer, as these extraembryonal tissues release ABA in response to far-red light and thereby inhibit germination (Lee et al., 2012; Yan et al., 2014). Therefore, we tested if the light inhibition of germination in Aethionema CYP seeds was also mediated by the seed coat and endosperm. Indeed, after mechanical removal of the extraembryonal tissue 24 hours after imbibition under light exposure, CYP and TUR seedling development was similar, and 100% of the seedlings grew normally, even under continuous light (Fig. 6A). These data indicate that, although Arabidopsis and Aethionema CYP seeds respond differently to light, the role of the seed coat and endosperm, and likely the involvement of ABA, appear similar.

**Fig. 6.**
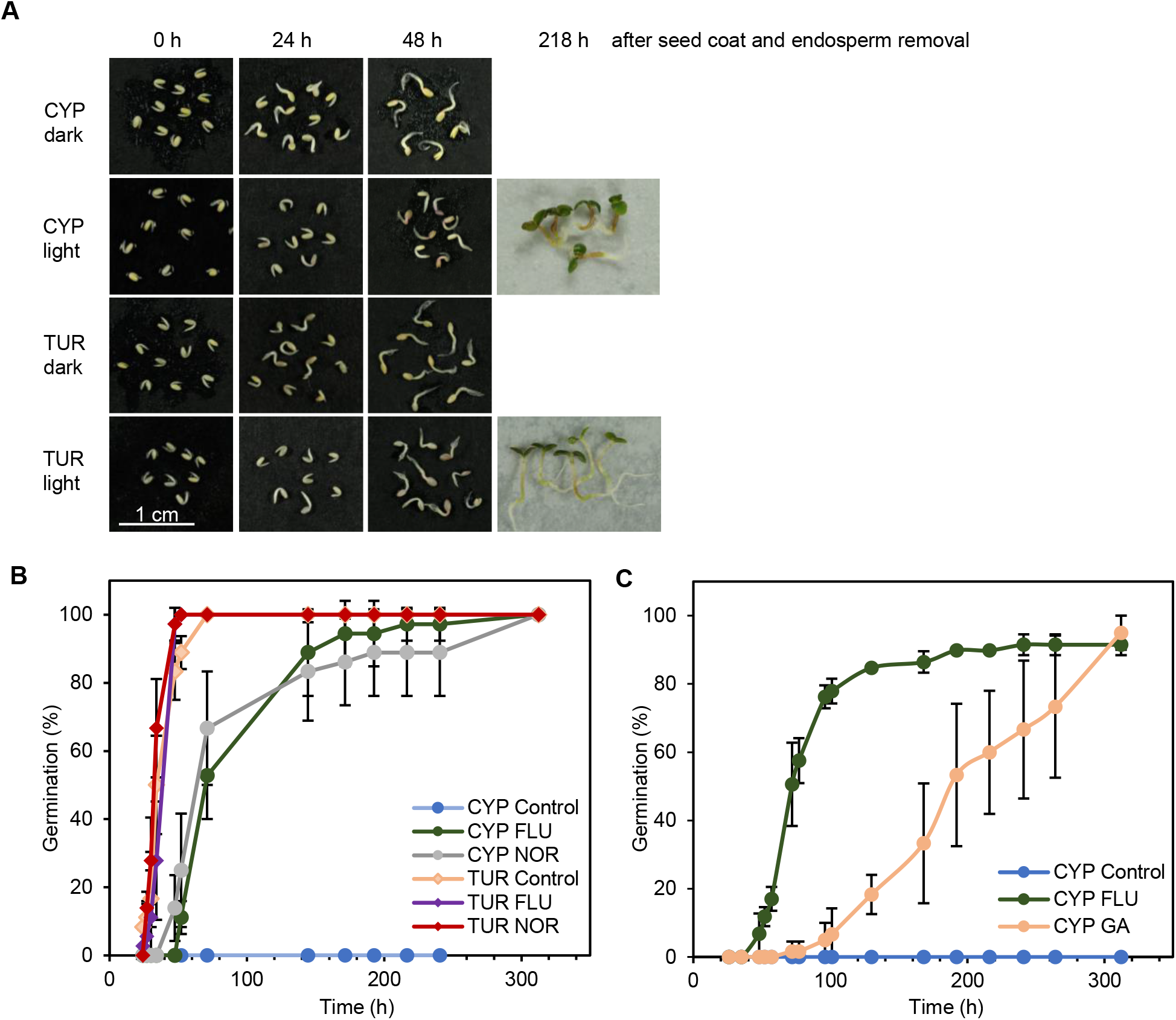
Rescue of CYP seed germination in light by interference with ABA and GA. **(A)** Development of seedlings in darkness or light after removal of the ABA-producing seed coat and endosperm. 0 hr indicates the status directly after seedling isolation. **(B)** Percentage of germination over time when seeds were kept on plates supplemented with ABA inhibitors 10 µM fluridone (FLU) or 10 µM norflurazon (NOR), or **(C)** external application of 10 µM GA_4+7_ (GA) or 0.01% DMSO as a control under continuous light. TUR (Turkey accession), CYP (Cyprus accession). Error bars represent standard deviation of three independent replicates.

Germination of light-requiring lettuce seeds in the dark or under far-red light could also be rescued by addition of norflurazon [4-chloro-5-methylamino-2-(3-trifluoromethylphenyl)pyridazin-3-one] (Widell et al., 1981). Fluridone [1-methyl-3-phenyl-5-(3-trifluoromethyl-phenyl)-4-(1H)-pyridinone] restored the seed germination of lettuce and other seeds at sub-optimal temperatures (Yoshioka et al., 1998; Debeaujon & Koornneef, 2000; Argyris et al., 2008). Both chemicals are inhibitors of carotenoid biosynthesis, which is required for *de novo* ABA synthesis (Bartels & Watson, 1978). We applied fluridone and norflurazon for both Aethionema accessions under continuous light exposure. Importantly, CYP seed germination was completely rescued (Fig. 6B), indicating that *de novo* ABA synthesis induced by light is an important component of the negative germination control in light-exposed CYP seeds.

Next, we tested if externally applied GA could overcome the light inhibition. Indeed, addition of 10 µM GA4+7 allowed CYP seeds to germinate under continuous light, although the germination was slower compared to fluridone (Fig. 6C). These data suggest that GA and ABA are indeed involved in the control of germination as in other plants. However, the signaling pathways downstream of light reception to the transcriptional control of these two key hormones must be antipodal to those in Arabidopsis.

## Discussion

### Ecological significance of light-regulated germination

The first observations on photoblastic differences were already reported more than a hundred years ago (Kinzel, 1913), and since then, several species have been described with a light-inhibited or light-neutral germination phenotype (Grime *et al*., 1981). Despite this, research in the past decades about the molecular control of seed germination focused nearly exclusively on the light-requiring germination of Arabidopsis. Here, we present an initial physiological and molecular characterization of light-neutral and light-inhibited germination in two accessions of Aethionema, another Brassicaceae species.

The light-requirement for seed germination is often considered a depth-sensing strategy associated with small seed size (Fenner and Thompson, 2005; Grime *et al*., 1981). As the light only penetrates a few millimeters into the soil, light dependence of small seeds ensures that the elongating hypocotyl will reach the surface before its resources are exhausted (Fenner and Thompson, 2005; Woolley and Stoller, 1978). Additionally, the light quality, sensed by the phytochrome photoreceptors as the ratio of red:far-red wavelengths, provides information about the leaf canopy, as leaves absorb more red than far-red light. Therefore, the minimum red:far-red ratio required for seed germination of a certain species may determine the optimal season when the competition is reduced, as it was shown for *Cirsium palustre* (Pons, 1984). In contrast, light-inhibited germination is plausible in species originating from open, arid, or semi-arid habitats where high light intensity is likely coupled with drought conditions, which could be unfavorable for young seedlings (Lai *et al*., 2016; Thanos *et al*., 1991). These conditions occur at many original locations of *Aethionema arabicum*. Our data indicate that the light-inhibitable germination of the CYP accession might be a photoperiod-sensing mechanism, as the seeds germinate well under short day but not under long day conditions.

The short daylength corresponds to the early spring days, when Aethionema germinates in its natural habitat. As the average lifespan of Aethionema is around 4 months, early germination is necessary to complete the life cycle and seed production before the dry and warm season. Similar to Aethionema CYP, seeds of the light-inhibited garden variant of *Nemophila insignis* germinate preferentially in short days (Black and Wareing, 1960; Chen, 1968). Seasonal adaptation of germination via opposite photoperiod sensitivity was described for arctic tundra species: these are inhibited by short days and prefer to germinate under long days, corresponding to the short summer season in Alaska (Densmore, 1997). Despite these few known examples, photoperiod dependence of seed germination is likely more common for plants in habitats where optimal timing of germination is crucial.

Germination in most of the examined Aethionema species was at least partially inhibited by light, whereas the Turkish accession of Aethionema (TUR) and one *Ae. heterocarpum* accession germinated independently of light. The occurrence of both phenotypes among close relatives in the Aethionemeae indicates that it is likely an adaptive trait that appeared more than once during evolution, although the exact environmental cues that favor light-neutral germination is unknown. With the limited number of Aethionema accessions available that have been propagated under controlled conditions, it is too early to conclude which of the phenotypes is ancestral. Based on the habitats of most Aethionema species and the transcriptome changes of key regulatory genes in the same direction but to different degrees in TUR and CYP, it is tempting to speculate that light-inhibited germination is the ancestral mechanism that has been desensitized in some instances. Collection and amplification of seed material and analysis of further Aethionema species and accessions, for which detailed phylogenetic data are available (Mohammadin *et al*., 2017a), are expected to provide an answer to this question.

### Light-dependent versus light-inhibited germination control

The species for which light inhibition of seed germination was previously described (Botha and Small, 1988; Chen, 1970; Chen, 1968; Thanos *et al*., 1991) are phylogenetically distant from the model plants lettuce and Arabidopsis, which show light-dependent germination. Some molecular aspects of light-inhibited germination have been investigated for monocotyledonous plants (Barrero *et al*., 2012; Hoang *et al*., 2014), in both cases clearly restricted to blue light. The comparison within the triangle of closely related species with three germination phenotypes, light-requiring Arabidopsis, light-neutral Aethionema TUR, and Aethionema CYP that is inhibited by the full spectrum of visible light, may allow us to understand the mechanistic and evolutionary divergence of the light controlled signaling network inducing germination. Previous studies in numerous species (Finch-Savage and Leubner-Metzger, 2006) along with the data presented here leave no doubt of the central role of ABA and GA in the inhibition or stimulation of germination, respectively. The positive, essential stimulus of light in Arabidopsis and its negative, blocking role in Aethionema CYP are expected to reflect a fundamental and qualitative difference between light reception and hormonal control. Although many other factors, including other hormones, are known to modulate germination (Argyris *et al*., 2008; Linkies and Leubner-Metzger, 2012; Meng *et al*., 2016), the antipodal transcript level changes upon light exposure for some of the same ABA/GA key regulatory components as in Arabidopsis indicate a major source of the difference in this signaling pathway (Fig. 7). Among these are *AearSOM*, *AearABI3*, *AearABI5*, *AearABA1*, *AearNCED6*, *AearGA3ox1*, and *AearGA3ox2*. However, the expression of many other genes responds similarly in Aethionema and Arabidopsis, for example *RGA*, *GAI*, *DAG1*, *CYP707A2*, and *JMJ20*, indicating that the light response is partially conserved (Fig. 7). In Arabidopsis, PIL5/PIF1, a key regulator in the light-induced transcriptional cascade, undergoes a rapid protein degradation (Shen *et al*., 2008). As the antibody against the Arabidopsis PIL5/PIF1 did not recognize the Aethionema protein from the orthologous gene (*AA33G00286*), we were unable to test its light-responsive protein degradation in Aethionema seeds. However, it is remarkable that the direct downstream target genes (*DAG1*, *SOM*, *DELLAs*, *ABI3*, *ABI5*) are either up- or downregulated in Aethionema, whereas their transcriptional repression by light is rather uniform in Arabidopsis (Fig. 7). A study based on chromatin immunoprecipitation in Arabidopsis with the PIL5/PIF1 antibody and microarray data revealed 166 genes that are under the direct control of PIL5/PIF1 (Oh *et al*., 2009). Those Aethionema orthologues of the direct PIL5 target genes that could be identified (132 out of the 166) were found to have relative stable and light-independent expression in Aethionema seeds; only 8 and 7 genes in TUR and CYP, respectively, showed more than 2-fold changes in either direction (Dataset **S6**). Therefore, one possible divergence between Arabidopsis and Aethionema might be the regulation of PIL5/PIF1 protein activity or stabilization. The dark and far-red germination of Arabidopsis *pil5* mutant seeds further indicates that PIL5/PIF1 is the most upstream element in the network that is possibly associated with germination in the dark.

**Fig. 7.**
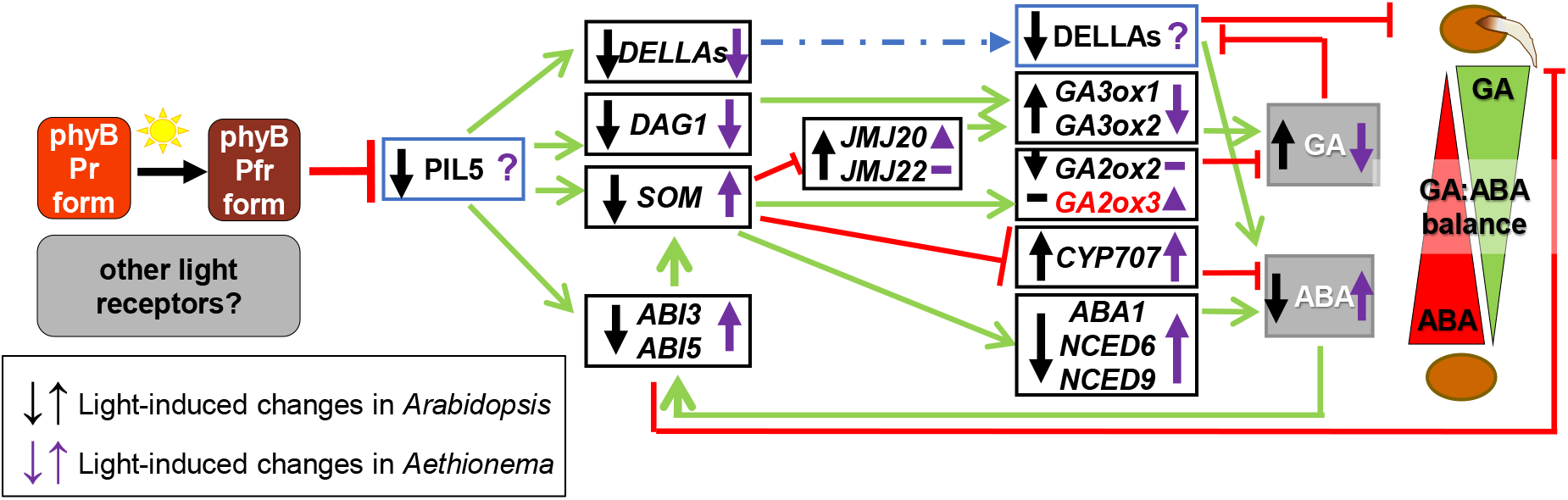
Antipodal transcriptional changes in Arabidopsis and Aethionema of key hormone regulatory genes. Summary of light-regulated transcript level changes in seeds, based on published Arabidopsis data and Aethionema data presented in this study. Blue boxes with straight letters indicate proteins whose stability is regulated by light. Black boxes with italic letters indicate genes whose expression is light regulated. The changes of expression upon light exposure are indicated by black (Arabidopsis) and purple (Aethionema) arrows, respectively. The role of other photoreceptors in Aethionema seed germination is not yet known. As marked with purple question marks, the protein stability of PIL5 and DELLAs are unknown in Aethionema.

### Light-independent versus light-inhibited germination control

Our transcriptome analysis identified 87 genes which are light-responsive and differentially regulated between the TUR and CYP accessions. Nevertheless, almost all 87 genes showed the same direction of transcript level change in both accessions, but the response was clearly more pronounced in CYP seeds. One possible explanation for this observation might be a reduced sensitivity of TUR seeds germination to light. In Arabidopsis, screens for light hyposensitive mutants often identified *PHYB* or *PHYA* mutations, indicating that the primary reason for hyposensitive response may be variations within the phytochrome protein. Like Arabidopsis, there are 5 phytochromes in Aethionema, which are highly conserved (Fig. **S4A**). The protein sequences of PHYB, PHYC, and PHYD are identical in both accessions, while the PHYA and PHYE proteins harbor a few missense SNPs (Fig. **S4B-F**). However, the second, light-inhibited accession from Turkey (KM2397) shares most of the same SNPs with the light-neutral TUR accession. It is therefore unlikely that allelic variations of phytochromes are responsible for the different responses (Fig. **S4B-F**). Similarly, there are no SNPs in the PIL5/PIF1 coding sequences that would cause non-synonymous amino acid changes and that diverge between the light-neutral TUR and the two light-inhibited accessions (Fig. **S5**). A detailed analysis about the phytochrome actions may help to understand and identify the photoreceptors involved in the light-inhibited germination.

As the qRT-PCR data indicated, many genes of the light regulated network are differentially expressed either slightly or strongly in TUR and CYP seeds under dark and light conditions, resulting in substantial differences in *AearGA3ox1* and *AearGA2ox2* expression and lower GA_4_ hormone level in CYP seeds under light. The differential ABA:GA ratio is likely determined by more than one upstream event early in the transcriptional cascade. Our data show that the *PAR1* and *PAR2* genes are significantly differentially expressed in TUR and CYP seeds under dark and light conditions. *PAR1* and *PAR2* both encode bHLH proteins, but do not possess DNA binding activity and are involved in shade avoidance response (Roig-Villanova *et al*., 2007; Wray *et al*., 2003). Although their precise mechanism and role in seed germination is still unknown, it has been speculated that they form heterodimers with other bHLH proteins, like PIFs, and modulate their activity as transcriptional cofactors (Roig-Villanova *et al*., 2007). Therefore, the differential expression of *AearPAR1* and *AearPAR2* in TUR and CYP seeds might play a role in the regulation of PIL5/PIF1 activity, influencing the network downstream of PIL5/PIF1.

While our data did not identify a unique point of divergence in the molecular control of light over seed germination between the investigated Aethionema accessions, the identified natural variation within the genus and the phylogenetic relationship with the conversely responding Arabidopsis provide great opportunities to elucidate the mechanism of an ecologically important, but under-investigated trait. Based on the current evidence, the basic components seem to be conserved but connected in a different cascade of events. Growth conditions, genome size, and generation time of Aethionema are similar to Arabidopsis, allowing for future forward mutant screens and genetic association studies.

## Acknowledgements

This work is part of the ERA-CAPS “SeedAdapt” consortium project (https://www.seedadapt.eu/). We thank all members for fruitful cooperation and discussion. We are especially grateful for Eric M. Schranz, Klaus Mummenhof, and Setareh Mohammadin to provide various Aethionema seed stocks. We also thank the staff at the Vienna BioCenter Core Facilities GmbH (VBCF), member of Vienna BioCenter (VBC), Austria, especially the Plant Sciences Facility for growth of the plants and the wavelength-specific light experiments, and the Next Generation Sequencing Facility for generating the RNA-seq data. We thank Nicole Lettner and Sarhan Khalil for technical support. We further acknowledge critical reading of the manuscript by J. Matthew Watson, Frederic Berger and Peter Hedden. The work was funded by the Austrian Science Fund (FWF) to O.M.S. (FWF I1477), by the Deutsche Forschungsgemeinschaft (DFG) to S.A.R. (RE1697/8-1), by the Biotechnology and Biological Sciences Research Council (BBSRC) to G.L.-M. (BB/M00192X/1), by a Natural Environment Research Council (NERC) Doctoral Training Grant to W.A. (NE/L002485/1), and the Czech Ministry of Education Youth and Sports grant LO1204 from the National Program of Sustainability I and Agricultural Research to M.S.

## Author contributions

Z.M., K.G., M.S., S.A.R., G.L.-M., and O.M.S. planned and designed the research; Z.M., K.G., W.A., D.T., and V.T. performed experiments; Z.M., K.G., P.W., K.K.U., W.A., C.G., D.T., V.T., M.S., S.A.R., G.L.-M. and O.M.S. analyzed and interpreted the data; Z.M., G.L.-M., and O.M.S. wrote the article; all authors approved the submitted version.

The following **Supplementary Information** is available for this article:

**Fig. S1** Heatmap of all 87 genes light-regulated in *Aethionema arabicum* CYP seeds and differentially expressed in light-exposed TUR and CYP seeds based on RPKM (reads per kilobase of transcript per million mapped reads) values.

**Fig. S2** Identification of the Arabidopsis orthologue of Aethionema *AA18G00108* as *GA2ox3*. A) Gibberellin 2-oxidase family phylogeny based on protein sequences. (B) Synteny of *GA2ox3* position in the genome of Arabidopsis and Aethionema.

**Fig. S3** Accumulation of GA forms in *Aethionema arabicum* TUR and CYP seeds under dark and light conditions.

**Fig. S4** Identification and alignments of phytochromes in *Aethionema arabicum*. (A) Phylogenetic tree of phytochromes. (B-F) Phytochrome A, B, C, D, E protein alignments of three *Aethionema arabicum* accessions.

**Fig. S5** Alignment of PIL5/PIF1 protein sequence of three *Aethionema arabicum* accessions.

**Table S1** Information about geographic origin of *Aethionema arabicum* accessions.

**Table S2** List of primers used for quantitative RT-PCR analysis.

**Table S3** List of Aethionema accession numbers used for this study.

## Methods S1

**Dataset S1**. List of differentially expressed *Aethionema arabicum* genes in TUR Dark versus TUR Light.

**Dataset S2**. List of differentially expressed *Aethionema arabicum* genes in CYP Dark versus CYP Light.

**Dataset S3**. List of differentially expressed *Aethionema arabicum* genes in CYP Dark versus TUR Dark.

**Dataset S4**. List of differentially expressed *Aethionema arabicum* genes in CYP Light versus TUR Light.

**Dataset S5**. List of common differentially expressed *Aethionema arabicum* genes in CYP Light versus TUR Light and TUR Dark versus TUR Light.

**Dataset S6**. List of target genes of Arabidopsis PIL5/PIF1 and transcriptional changes of orthologues in the Aethionema experiments.

**Dataset S7**: List of plant species for which protein sequences were considered for phylogenetic tree constructions. 24

